# Comparative co-expression reveals a regulatory core shared by angiosperm and conifer roots under cold

**DOI:** 10.64898/2026.08.15.745010

**Authors:** Tuuli Aro, Elena M van Zalen, Alexander Vergara, Camilla Canovi, Vikash Kumar, Ellen Dimmen Chapple, Torgeir R. Hvidsten, Vaughan Hurry, Nathaniel R. Street

**Affiliations:** Umeå Plant Science Centre (UPSC), Department of Forest Genetics and Plant Physiology, Swedish University of Agricultural Sciences, 901 83 Umeå, Sweden; Umeå Plant Science Centre (UPSC), Department of Plant Physiology, Umeå University, 901 87 Umeå, Sweden; Instituto de Ciencias de la Ingeniería, Universidad de O’Higgins, Rancagua, Chile; Faculty of Chemistry, Biotechnology and Food Science, Norwegian University of Life Sciences, 1433 Ås, Norway; SciLifeLab, Department of Plant Physiology, Umeå University, 901 87 Umeå, Sweden

**Keywords:** boreal forest, cold acclimation, comparative co-expression, conserved gene regulation, gibberellin, gymnosperm, RNA sequencing, root transcriptome

## Abstract

• Climate change is reducing the boreal snowpack that insulates soils, exposing tree roots to more frequent freezing. The molecular cold response of roots is poorly characterised, and its conservation across the angiosperm–gymnosperm divide is unknown. We asked how much of the root cold response is shared among divergent boreal trees.

• We profiled fine-root transcriptomes of four boreal trees (*Picea abies*, *Pinus sylvestris*, *Betula pendula* and *Populus tremula*) and two *Arabidopsis thaliana* ecotypes during a ten-day 5 °C treatment by RNA sequencing, and used comparative co-expression to detect conserved regulation independently of response timing.

• Expressed genes were largely shared, but the genes differentially expressed, and the timing of their response, diverged between species. Despite this, a core of orthologues retained conserved co-expression neighbourhoods across more than 300 million years, enriched for growth regulation, metabolism and stress signalling, including a gibberellin-related metabolic component of the growth response.

• The root cold response is therefore species specific in identity and timing, yet underlain by a conserved co-expression core.

## Introduction

Anthropogenic warming is altering growth conditions for plants worldwide, and the boreal zone is experiencing some of the most rapid and extreme change (Pörtner et al. 2022; Rantanen et al. 2022). Rising mean temperatures are shortening the winter, and an increasing fraction of boreal precipitation is falling as rain rather than snow (Bintanja & Andry 2017; Douglas et al. 2020). Boreal plants cold acclimate, increasing their freezing tolerance in the autumn in response to shortening days and falling temperatures (Sakai & Larcher 1987; Körner 2016), but they also depend on stable winter conditions, in which continuous snow cover insulates the soil and understorey against air-temperature fluctuations (Vuosku et al. 2022). Warmer, wetter winters are expected to reduce the depth and persistence of the annual snowpack (Mellander et al. 2007; Blume-Werry et al. 2016), and reduced snow insulation is predicted to increase the severity and depth of soil frost and to delay spring thawing (Eurola 1975; Richardson et al. 2024), all of which heighten the vulnerability of root systems to freezing injury (Schwartz et al. 2006; Martz et al. 2016; Domisch et al. 2018; Richardson et al. 2024).

Roots are generally more susceptible to freezing injury than above-ground tissues (Stier et al. 2003), and fine roots, in particular, have limited capacity to acclimate relative to lignified roots (Ryyppö et al. 1998). Declining soil temperatures during winter and spring are therefore expected to damage fine roots and suppress root growth, impairing water and nutrient uptake (Repo et al. 2014; Repo et al. 2021; Li & Hoch 2025) and ultimately reducing whole-plant productivity (Mellander et al. 2006; Pan et al. 2011). Recent physiological work shows that low root-zone temperature progressively restricts water uptake and transport in temperate and boreal trees, with the magnitude of this restriction varying among species (Li & Hoch 2025). This vulnerability is of particular concern because boreal forests are a major component of the global forest carbon sink (Pan et al. 2011). Understanding how the roots of boreal trees respond to cold is therefore important for anticipating whether these forests can maintain their role in carbon sequestration under a changing climate.

To date, the genomics of abiotic stress has focused overwhelmingly on herbaceous model angiosperms, and it remains uncertain how well the underlying processes generalise across the plant kingdom (Meng et al. 2021; Kim et al. 2024). Cold acclimation requires the coordinated activation of multiple regulatory pathways that are initiated within minutes to hours of low-temperature exposure and can continue for days to weeks (Moliterni et al. 2015; Zhao et al. 2020; Zhao et al. 2021; Perez-Garcia et al. 2023; Ma et al. 2024). In angiosperms, a central axis of this response is the ICE1–CBF–COR pathway, in which the CBF/DREB1 transcription factors induce downstream cold-regulated genes (Cook et al. 2004; Park et al. 2015; Qian et al. 2024). Comparative studies are increasingly revealing the limits of this paradigm. Cross-species analyses in grasses (Schubert et al. 2019), Brassicaceae (Birkeland et al. 2020; Birkeland et al. 2022) and a broad sample of angiosperms (Wang et al. 2023; Andrew et al. 2025) show that orthologous genes frequently differ in their cold responses between species, although machine-learning classifiers trained on one species can nonetheless predict which orthologues respond to cold in another (Meng et al. 2021). At the same time, multi-omics analyses across deeply diverged rosids have identified a conserved, hierarchically organised cold-responsive regulatory network, indicating that conservation and divergence coexist within the same response (Guo et al. 2023); comparative analyses across angiosperms have since defined sets of conserved cold-responsive transcription-factor orthogroups, including WRKY and NAC modules with demonstrated function (Jia et al. 2026).

The growing availability of tree genome assemblies, including boreal species such as Norway spruce (*Picea abies*) and Scots pine (*Pinus sylvestris*) (Nystedt et al. 2013; Ahlgren Kalman et al. 2025), silver birch (*Betula pendula*) (Salojärvi et al. 2017) and aspen (*Populus tremula*) (Robinson et al. 2024), now makes comparative transcriptomic analysis of stress responses in trees feasible. Comparative genomics identifies conserved and lineage-specific gene families; coupling this with comparative analysis of expression identifies orthologues with conserved responses, and comparative co-expression analysis extends this to conserved regulatory relationships. A key strength of the co-expression approach is that conservation does not require that orthologues share the same expression level or dynamics, only that they share co-expressed neighbourhoods, allowing conserved regulation to be detected even when the timing of responses differs (Movahedi et al. 2011; Netotea et al. 2014; Ovens et al. 2021; Crow et al. 2022). Established tools for such analyses, including ComPlEx (Netotea et al. 2014) and PlaNet (Mutwil et al. 2011), integrate large-scale expression data with orthology, but their application has been largely restricted to angiosperms, owing to data availability and the fragmented annotation of gymnosperm genomes.

Previous comparative work between *Arabidopsis thaliana* (Arabidopsis) and Norway spruce established that the timing of the cold response differs between these species and that responses are tissue specific, with roots and needles deploying distinct programmes (Vergara et al. 2022); a comparable pattern of tissue-specific regulation has also been reported for drought responses in the same species (Haas et al. 2021). However, that work was confined to two species, and roots in general remain far less studied than above-ground tissues; no broad, multi-species comparative transcriptomic study of the cold response of tree roots has yet been conducted, and none has included gymnosperms alongside angiosperms in a common framework. Resolving the extent of shared versus divergent regulation in roots is a prerequisite both for transferring mechanistic knowledge between species and for designing targeted strategies to improve root performance under cold soils.

In this study we used one-year-old seedlings of four native boreal trees (the gymnosperms Norway spruce and Scots pine and the angiosperms birch and aspen) together with the Arabidopsis ecotypes Columbia (Col-0) and the northern Östhammar (Ost-0), chosen to span the latitudinal variation in freezing tolerance and CBF-independent cold regulation documented among Arabidopsis accessions (Zuther et al. 2012; Park et al. 2018), to assess the core cold response of boreal tree roots. We profiled fine-root transcriptomes by RNA sequencing over a ten-day exposure to 5 °C (a chilling treatment that elicits cold acclimation, the process underlying freezing tolerance, rather than freezing injury itself), and asked three questions: (i) how much of the differentially expressed cold response is shared among species; (ii) are genes with shared temporal response dynamics orthologous, and therefore represent conserved mechanisms; and (iii) can comparative co-expression recover conserved regulatory relationships that are obscured by the divergence and differing timing of differential expression. The comparative co-expression framework applied here was previously used to compare cold responses between Norway spruce and Arabidopsis (Vergara et al. 2022) and to reconstruct the regulation of wood formation across dicot and conifer trees (Rodriguez et al. 2026). Here we apply it to the root cold response across a broader sampling of six boreal species and ecotypes, extending the approach from a developmental programme to an environmental stress and, by including the herbaceous annual Arabidopsis alongside the perennial trees, enabling a life-history contrast. This addresses whether conserved co-expression is a property of developmental programmes alone or also extends to stress responses.

## Materials and methods

### Plant material, experimental design and sampling

One-year-old seedlings of Scots pine (*Pinus sylvestris* L.; seed provenance Sönnersta, 63° 28′ N), birch (*Betula pendula* Roth; provenance Sävar, 63° 54′ N) and Norway spruce (*Picea abies* (L.) H. Karst; provenance Lilla Istad, 56° 30′ N) were grown in 1-l pots at 18 °C/15 °C (light/dark) under an 18-h photoperiod. At the start of the experiment, seedlings were transferred to 5 °C, two hours into the photoperiod. Fine roots (<2 mm) were sampled at 0 h, 6 h, 24 h, 3 days and 10 days, rinsed in MilliQ water, blotted dry, frozen in liquid nitrogen and stored at −80 °C.

Aspen (*Populus tremula* L.; genotype SwAsp 76, central Sweden, Hälsingland, 61° 42′ N) was clonally propagated from tissue culture. Plantlets were established in 1-l boxes on Murashige and Skoog medium with gelrite (Duchefa Biochemie, the Netherlands) and amplified for four months at 22 °C/18 °C (light/dark) under an 18-h photoperiod. Plantlets were then transplanted to 1-l pots, allowed to acclimate and grow for 40 days, and subjected to the same cold treatment and sampling scheme as the other trees.

Seeds of Arabidopsis (*Arabidopsis thaliana* (L.) Heynh.) Col-0 (52° 42′ N) and Ost-0 (60° 15′ N) were stratified for 48 h at 5 °C in darkness, sown into individual pots and grown under a 16-h photoperiod at 23 °C. After 18 days, plants were transferred to 5 °C two hours into the photoperiod, and roots were sampled at 0 h, 6 h, 24 h, 3 days and 10 days, rinsed in 5 °C water, blotted dry and flash-frozen as above.

### RNA extraction and sequencing

Total RNA was isolated using a combination of the Qiagen RNeasy Plant Mini Kit (Qiagen) and the cetyltrimethylammonium bromide (CTAB) method of Chang et al. (1993) with minor modifications. Frozen samples were ground twice for 30 s at 25 Hz in a TissueLyser II (Qiagen), with cooling in liquid nitrogen between grinds. Warm (65 °C) extraction buffer (700 µl) was added and the samples were extracted twice with chloroform:isoamyl alcohol (24:1), with centrifugation at 10,000 rpm for 10 min. Nucleic acids were precipitated in ethanol, pelleted by centrifugation, resuspended in RLT buffer and processed through the RNeasy column protocol, including on-column DNase digestion (RNase-Free DNase Set, Qiagen). RNA was quantified on a NanoDrop 1000 (Thermo Fisher Scientific) and assessed for integrity on a Bioanalyzer 2100 with the RNA Nano 6000 kit (Agilent). Samples with an RNA integrity number above 8 were sequenced on the Illumina HiSeq 2500 platform (Novogene, UK).

### Read processing and quantification

Raw reads were quality-assessed with FastQC v0.11.9 (Andrews 2010), depleted of ribosomal RNA with SortMeRNA v4.3.4 (Kopylova et al. 2012) and adapter- and quality-trimmed with Trimmomatic v0.39 (Bolger et al. 2014), followed by a second round of FastQC. Reads from each species were quantified against the corresponding genome/transcriptome assembly with Salmon v1.8.0 (Patro et al. 2017): *A. thaliana* (AtRTD3 v1.6.0; Zhang et al. 2022), *P. tremula* (v2.2; Robinson et al. 2024), *B. pendula* (v1.4; Salojärvi et al. 2017), *P. abies* (v2.0) and *P. sylvestris* (v1.0) (both Ahlgren Kalman et al. 2025). Statistical analysis was performed in R v4.3.1 (R Core Team 2024). Transcript-level estimates were imported and summarised to the gene level with tximport v1.28.0 (Soneson et al. 2015). Counts were used directly for differential-expression testing, whereas a variance-stabilising transformation (VST; DESeq2 v1.40.2; Love et al. 2014) was applied to generate the normalised values used for sample quality control, distance-based clustering and co-expression analysis. Biological replicate quality was assessed by principal component analysis and hierarchical clustering of VST values.

### Differential expression and functional enrichment

Biological replication per species (control/6 h/24 h/3 d/10 d) was: Col-0 5/5/5/5/5; Ost-0 5/5/5/5/5; aspen 5/5/5/5/5; birch 5/5/5/5/5; Norway spruce 3/3/3/3/3; Scots pine 2/4/5/5/2 (25, 25, 25, 25, 15 and 18 samples per species, each forming that species’ co-expression network). Differential expression was tested in DESeq2 (Love et al. 2014), which models counts with a negative-binomial generalised linear model; differential expression used a single-factor design in which one factor (Condition) encoded the untreated control and each cold timepoint (6 h, 24 h, 3 d and 10 d at 5 °C); the control formed a single common reference group against which each cold timepoint was contrasted. Genes were considered differentially expressed (DEGs) at an adjusted *P* ≤ 0.05 and an absolute log2 fold change ≥ 1 (a two-fold change). The fold-change criterion was applied as a post-hoc filter to the standard Wald test (whose null is no change) rather than through a fold-change-thresholded null, and so does not itself provide formal false-discovery-rate control for the composite hypothesis of a two-fold change. Orthogroups were inferred as described in Rodriguez et al. (2026), using OrthoFinder v2.5.2 (Emms & Kelly 2019) applied to the longest protein-coding sequence per gene across 27 plant species with a species tree rooted using TimeTree (Kumar et al. 2022); hierarchical orthogroups (HOGs) were used as the unit of cross-species comparison. Because orthogroups can contain multiple genes and gene copy number can vary among species, orthogroup counts do not correspond directly to gene counts. Gene Ontology (GO) enrichment was performed with topGO using Fisher’s exact test (classic algorithm) with Benjamini–Hochberg correction (Alexa et al. 2006), using annotation from PlantGenIE (Sundell et al. 2015).

### Cross-species superclusters

To identify groups of genes with shared temporal response dynamics across species, we focused first on transcription factors (TFs) that were differentially expressed in response to cold (TF-DEGs). We seeded the analysis with TFs, rather than with all DEGs, because as upstream regulators their cold-responsive dynamics are expected to define the temporal structure of the regulatory programme more sharply than the larger and noisier set of downstream structural genes; correlated non-TF DEGs were added subsequently so that the analysis captured both candidate regulators and their putative targets. For each species, TF-DEG expression profiles were grouped by hierarchical agglomerative clustering (Ward’s method) into six clusters with synchronised cold-responsive dynamics. Pairwise correlations between all clusters across species were then used to group the clusters manually (Pearson r ≥ 0.7) into twelve cross-species superclusters (SCs), each representing a distinct temporal response pattern (Fig. S1). Some SCs contained clusters from multiple species and others only single-species clusters. To capture co-regulated genes excluded by the initial clustering and to incorporate putative downstream targets, each SC was populated with additional TF-DEGs and non-TF DEGs whose expression profiles correlated with the SC mean (Pearson *r* ≥ 0.7). Because genes are recruited by their correlation with the supercluster mean, this step defines supercluster membership rather than testing it; we therefore treat the superclusters as descriptive, and the conclusion drawn from them — that co-timed genes are largely not orthologous — does not depend on this recruitment. Subsequent analyses focused on the four SCs (SC_I–SC_IV) that contained TF-DEGs and more than 20 correlated DEGs in all species. These four superclusters were renumbered SC_I– SC_IV for presentation and correspond, respectively, to superclusters SC6, SC2, SC3 and SC1 of the twelve shown in Fig. S1.

### Comparative co-expression analysis

Conserved co-expression relationships were identified with the Comparative analysis of Plant co-Expression (ComPlEx) framework (Netotea et al. 2014), following Rodriguez et al. (2026). For each species, genes were retained for network construction if they had non-zero variance, non-zero expression in at least three samples, and a mean expression above 1.0 on the variance-stabilised scale (shifted so that each gene’s minimum is zero); a co-expression network was then constructed from Pearson correlations. Mutual ranking was applied to improve robustness, and networks were trimmed to retain the strongest 3% of possible edges (density = 0.03); this density was fixed a priori, consistent with the ComPlEx framework (Netotea et al. 2014) as applied by Rodriguez et al. (2026), rather than tuned to these data. For each orthologue pair, a hypergeometric test assessed in both directions

whether co-expression neighbourhoods overlapped significantly between species, with Benjamini–Hochberg correction for multiple testing; throughout, “bidirectional” denotes that the larger of the two directional adjusted P values was required to fall below the threshold. Orthologue pairs with a bidirectional adjusted *P* < 0.1 were termed co-expressologs, and sets of orthologues (containing one gene per species) classified as co-expressologs in all pairwise species comparisons were termed cliques, following the cross-species comparative co-expression approach introduced by Rodriguez et al. 2026. Because orthogroups can contain several genes per species, each species was represented within a clique by a single gene; 110 of the 116 conserved-core orthogroups yielded more than one complete clique (median of 26 complete cliques per orthogroup, across all 116), and a single representative complete clique per orthogroup was used for the one-gene-per-species visualisation (Figs 5 and 6). The bidirectional test was considered significant only when both directions passed the threshold (equivalently, when the larger of the two adjusted P-values was below 0.1); because a conserved-core clique requires this reciprocal co-expressolog relationship to hold simultaneously across all 15 pairwise comparisons, the effective stringency at the clique level substantially exceeds the per-comparison threshold. To test whether the number of orthogroups conserved across all pairwise comparisons exceeded chance, we performed a permutation test in which co-expressolog assignments were randomised within each pairwise comparison, preserving the number of co-expressologs per comparison, and the number of orthogroups recovered as co-expressologs across all comparisons was recorded over 5,000 permutations.

As an independent check that the conserved-core clusters represent coherent co-regulation, we asked whether the Arabidopsis genes of the conserved core were more co-expressed within clusters than between them. We took co-expression links between Arabidopsis genes from the co-expression channel of STRING v12 (Szklarczyk et al. 2023; channel subscore ≥ 400), mapping gene identifiers from TAIR loci with the STRING alias table. We then restricted these links to the conserved-core genes and counted those falling within a clique cluster (Fig. 5). Finally, we compared this count against 5,000 permutations that randomly reassigned the same genes to clusters of the observed sizes.

## Results

### A largely shared transcriptome underlies a divergent cold response

The roots of the four boreal trees and the two Arabidopsis ecotypes expressed a broadly similar set of genes under both control and cold conditions, sharing 6,502 orthogroups (Fig. 1A) that accounted for approximately half of the expressed transcriptome in each species (Fig. 1B). The two conifers shared a large set of conifer-specific orthogroups and also had many unique orthogroups, consistent with their higher total gene number, the extensive lineage-specific gene-family expansion and gene duplication that characterise conifer genomes (Ahlgren Kalman et al. 2025), and the deep divergence both between gymnosperm and angiosperm lineages (more than 300 million years; Kumar et al. 2022) and between the two conifers themselves (∼130 million years; Ahlgren Kalman et al. 2025).

**Fig. 1.**
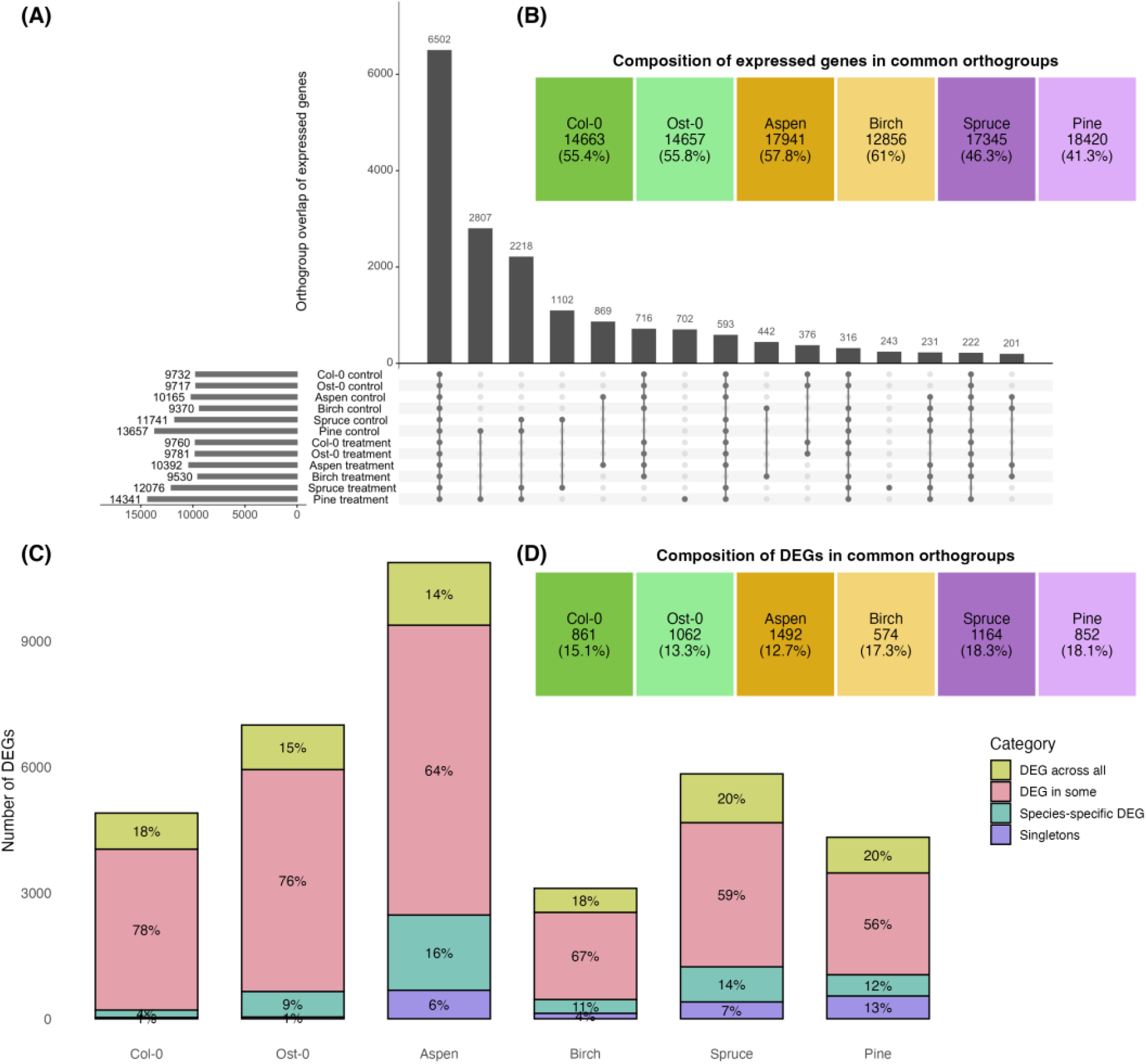
Orthogroup overlap of expressed genes and differentially expressed genes. (A) UpSet plot of orthogroups across all expressed genes. Horizontal bars give the total number of orthogroups in each set; vertical bars give the size of each intersection. (B) Gene number per species and the percentage of expressed genes lying in orthogroups expressed in common across all species/ecotypes. (C) Composition of differentially expressed gene (DEG) orthogroups: “DEG across all”, every species has a DEG in the orthogroup; “DEG in some”, more than one but not all species; “species-specific DEG”, a DEG in a single species while other species’ orthologues are not DEGs; “singletons”, a DEG with no orthologue in any other species. (D) Proportion of each species’ DEGs lying within the common (“DEG across all”) set.

Against the backdrop of shared gene content, the cold response itself diverged substantially between species. The largest category of cold-responsive DEGs had an orthologue that was also differentially expressed in at least one other species (Fig. 1C), yet only 270 DEG-containing orthogroups were shared across all six species/ecotypes. This common set represented only ∼4–8% of the commonly expressed genes and ∼12.7–18.3% of all DEGs in each species (Fig. 1D). Resolved among orthogroup-mapped DEGs (a smaller denominator than in panel D), 14–20% of DEGs had an orthologue that was differentially expressed in all species, 56–78% had an orthologue differentially expressed in at least one other species, 4– 16% had an orthologue present in another species but differentially expressed only in the focal species, and 1–13% were species-specific singletons (Fig. 1C). As the underlying gene content is largely shared, this divergence reflects differences in how a common repertoire of genes is regulated rather than differences in gene usage, with gene duplication within some orthogroups giving rise to copies that differ in their cold regulation. In addition to species-specific responses, Col-0 and Ost-0 each had ecotype-specific DEGs, indicating that divergence of the cold-stress transcriptome operates even between ecotypes of a single species. Nonetheless, the 270 commonly responding orthogroups were significantly enriched for GO terms including “response to oxygen-containing compound” and multiple categories associated with abiotic stress and hormonal signalling (Table S1), identifying a core of cold-responsive functions shared among species separated by hundreds of millions of years.

### Genes with shared response timing are largely not orthologous

Across species, roots differed markedly in the timing of differential expression over the ten-day treatment (Fig. 2). The fastest responses occurred in the Arabidopsis ecotypes Col-0 and Ost-0, whereas all tree species showed slower induction, and the two Arabidopsis ecotypes themselves differed in timing. This raised the question of whether any temporally coherent response was conserved at the level of orthologous genes. Clustering of cold-responsive TFs across species yielded four superclusters with characteristic temporal profiles (Fig. 3A; Fig. S2; Table S4). Three superclusters (SC_I, SC_II, SC_III) showed early induction with distinct kinetics: a transient peak at 6 h (SC_I), induction to 24 h followed by decline (SC_II), and induction within 24 h that was then sustained (SC_III). By contrast, SC_IV showed continuous up-regulation and contained the most DEGs of any supercluster (Fig. 3A, 3B). SC_I was dominated by Arabidopsis and contained a high proportion (62%) of Arabidopsis-unique orthogroups (Fig. 3C), whereas SC_II and SC_III were more evenly distributed among species. Critically, orthogroup overlap among species was low within every supercluster, for both TFs and non-TF genes (Fig. 3C). Thus, although genes following common temporal trajectories could be found in all species, those genes were generally not orthologous, so shared response timing alone does not indicate conserved regulation of the same genes, and low orthogroup overlap may also reflect the orthogroup-level analysis or incomplete conifer annotation.

**Fig. 2.**
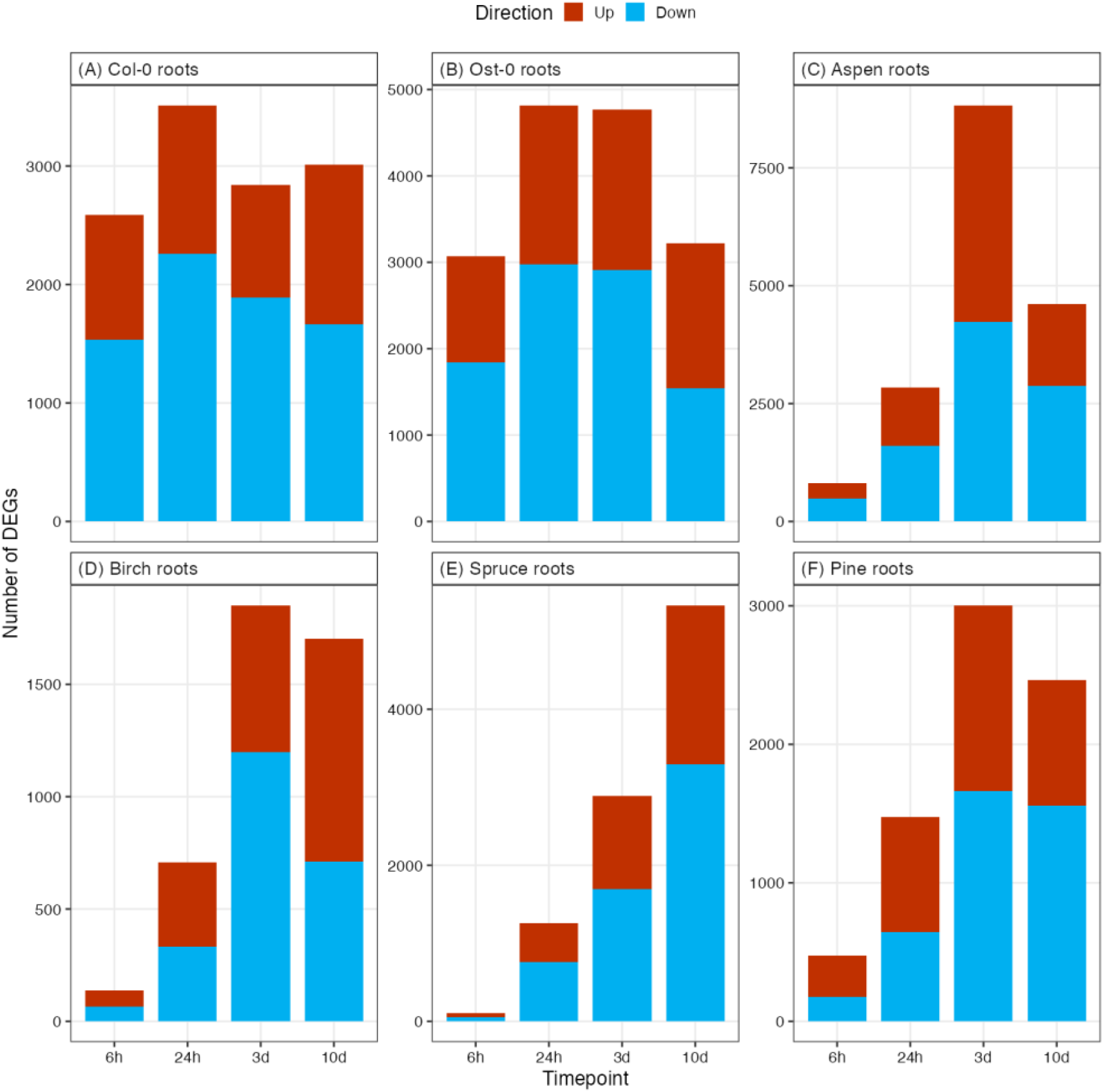
Numbers of up- and down-regulated differentially expressed genes (DEGs) during cold treatment (5 °C) at 6 h, 24 h, 3 d and 10 d in (A) *Arabidopsis* Col-0, (B) *Arabidopsis* Ost-0, (C) aspen, (D) birch, (E) Norway spruce and (F) Scots pine fine roots.

**Fig. 3.**
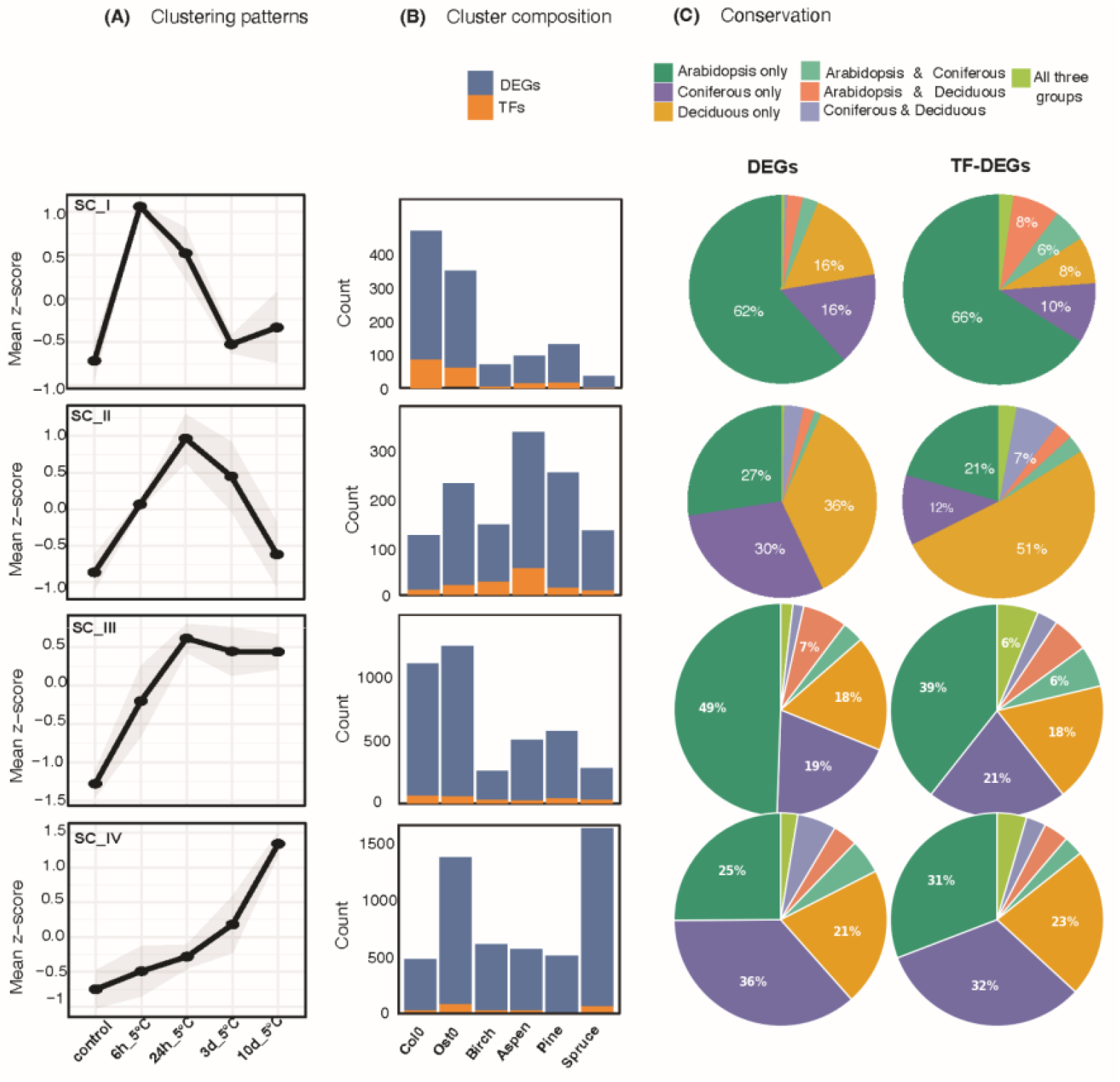
Supercluster (SC) characterisation. (A) The four SCs (I–IV) showing characteristic cold-response timing across species; points give the mean and shading the standard deviation of *Z*-score-normalised expression across species. (B) SC composition in terms of DEGs and transcription factors (TFs) per species. (C) Proportion of orthologous relationships among SC members (TF-DEGs and DEGs) across species, shown as percentages.

### Comparative co-expression recovers conserved and lineage-specific regulation

The divergent timing of cold responses across species (low overlap among superclusters), indicates extensive divergence in the differential-expression response of roots to cold. As conserved regulation can persist even when expression dynamics differ, we next applied comparative co-expression analysis, which identifies orthologues with conserved co-expression neighbourhoods (co-expressologs) independently of their response timing (Netotea et al. 2014). This recovered a large set of co-expressologs shared across all pairwise comparisons, irrespective of whether the genes were classified as differentially expressed (Fig. S3).

Among pairwise comparisons (Fig. S4), the most co-expressologs were shared between the two Arabidopsis ecotypes (1,404), followed by the two conifers (716) and the two broadleaf trees (291). Guided by these within-lineage similarities, we grouped the comparisons into three sets, Arabidopsis (A), broadleaf trees (B) and conifer trees (C), and examined their overlaps (Fig. 4A). A substantial shared set of 1,277 orthogroups with conserved co-expression was shared across all three sets (ABC), but the analysis also revealed conservation that was specific to different sets of species: The largest single set represented conserved co-expression within Arabidopsis ecotypes (A, 2,857 orthogroups), and a further 1,977 orthogroups were shared specifically between Arabidopsis and the broadleaf trees (AB). By contrast, the raw Arabidopsis–conifer overlap (AC, 639) exceeded the broadleaf– conifer overlap (BC, 133). This apparent phylogenetic paradox reverses once set size is taken into account: 36% of the broadleaf set (BC + ABC = 1,410 of 3,924 orthogroups) was conifer-conserved versus 28% of the Arabidopsis set (AC + ABC = 1,916 of 6,750), so the broadleaf set was 1.27-fold more conifer-conserved than the Arabidopsis set — consistent with shared tree biology. The larger raw Arabidopsis–conifer overlap therefore reflects the larger Arabidopsis co-expressolog set, itself likely a consequence of the more complete Arabidopsis annotation, rather than greater conservation (annotation asymmetry is considered further in the Limitations). The shared sets (ABC, AB and BC) were significantly enriched for GO terms, whereas the species-exclusive sets (A, B and C) showed little or no enrichment (Fig. 4B); the broadleaf- and conifer-exclusive sets (B and C) contained relatively few genes, which most likely explains their limited enrichment, whereas the Arabidopsis-exclusive set (A) was the largest region yet showed almost no enrichment. The shared core (ABC) was enriched for “response to stress”, “response to hormone” and “response to endogenous stimulus”, consistent with a deeply conserved component of the cold response (Table S2). The broadleaf–conifer (BC) set was enriched for “response to abscisic acid”, “defense response” and “gibberellin biosynthetic process” (Table S2), identifying responses shared among the trees but not Arabidopsis.

**Fig. 4.**
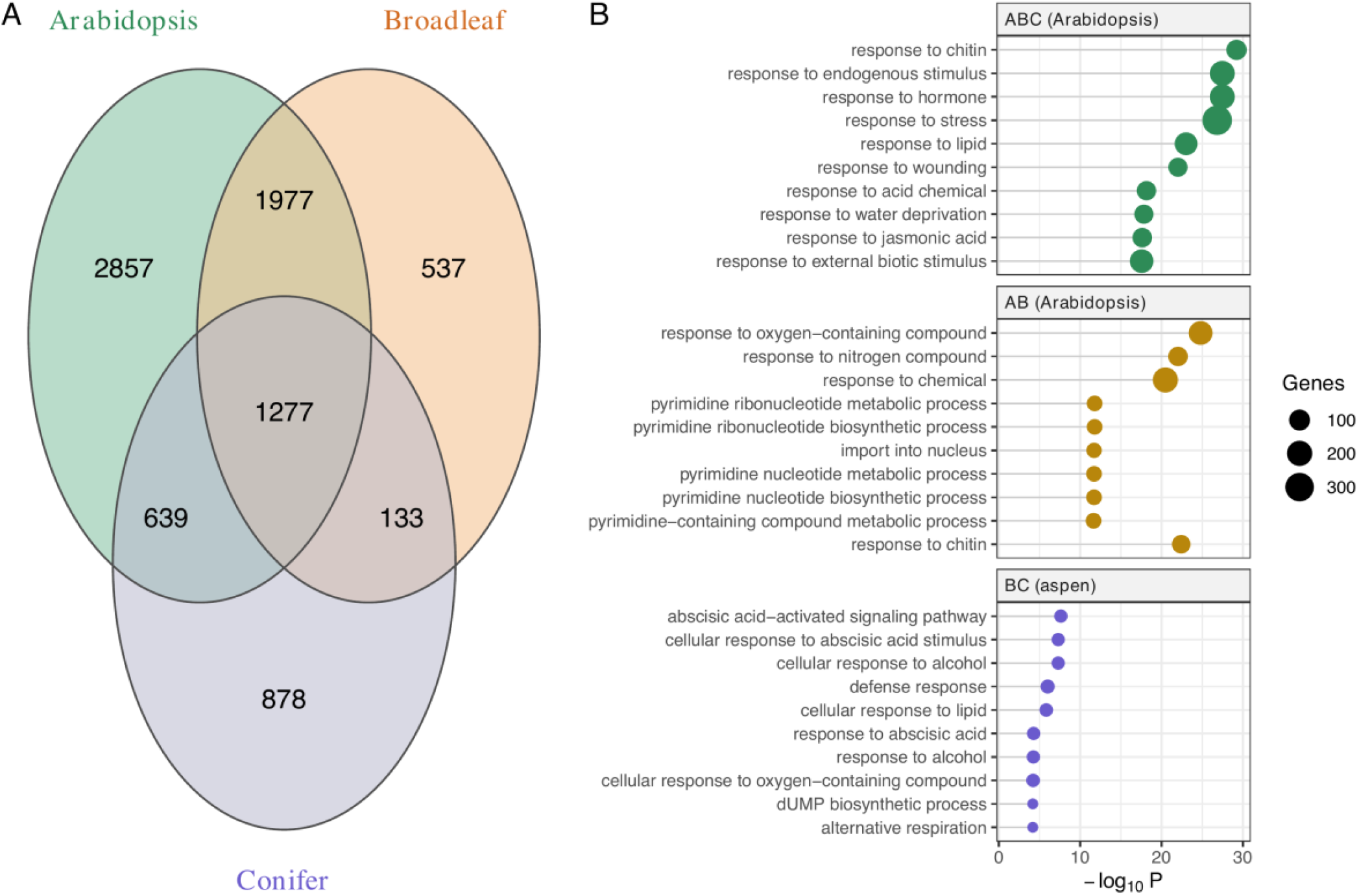
(A) Venn diagram of orthogroup overlap among co-expressologs of the within- *Arabidopsis* (A), within-broadleaf (B) and within-conifer (C) comparisons. (B) The ten most significantly enriched biological-process GO terms for the shared sets (ABC, AB, BC); point position is −log10 *P* (topGO classic Fisher, Benjamini–Hochberg padj < 0.05) and point size the number of genes. For sets including *Arabidopsis* the Col-0 annotation was used, otherwise the aspen (broadleaf) annotation. The species-exclusive sets (A, B) and the conifer-only set (C) showed little or no enrichment and are not shown.

### A conserved regulatory core spanning angiosperms and gymnosperms

To define the most strongly conserved component of the response, we identified 256 orthogroups whose co-expressolog pairs were present across all pairwise comparisons; of these, 116 formed fully interconnected cliques, containing one gene from each species, where every pair was a co-expressolog. These cliques represent cold-responsive regulatory relationships conserved across lineages separated by more than 300 million years (Kumar et al. 2022). Across 5,000 permutations that randomised co-expressolog assignments independently within each pairwise comparison, breaking the correspondence between comparisons, no orthogroup was recovered as a co-expressolog in all 15 comparisons (P < 0.0002; Fig. S5a). Because each clique selects a single corresponding gene per species, it provides a 1-to-1 mapping for clustering and visualising expression profiles directly across species. Genes within the cliques fell into four expression clusters (Fig. 5). Cluster 1 (50 orthogroups) was enriched for cell-division processes (“cell population proliferation”, “cytokinesis by cell plate formation”; enriched most strongly in the Arabidopsis ecotypes, Table S3) and showed a striking directional contrast: these genes were induced by cold in the two Arabidopsis ecotypes but down-regulated in birch, spruce and (by day 10) Scots pine, with aspen intermediate. This opposing direction could indicate maintenance of root growth in the herbaceous annual versus growth cessation in the perennial trees. Cluster 2 (30 orthogroups) was enriched for metabolic processes (“protein metabolic process” across most species and “RNA methylation” in the Arabidopsis ecotypes) and was induced in the Arabidopsis ecotypes, aspen and the two conifers, with differing timing (by 24 h in Arabidopsis, more diffusely in aspen, and only by day 10 in the conifers); birch was the exception, where these genes were repressed throughout the time course. Cluster 3 (20 orthogroups) was enriched for signalling and stress terms (“intracellular signal transduction”, “response to stress”, “response to abiotic stimulus”) and showed enrichment only in the two Arabidopsis ecotypes, with no significant terms in any tree species for this cluster (Table S3); Cluster 4 (16 orthogroups) showed no significant enrichment. The largest conserved component therefore concerns the regulation of root growth, indicating that adjusting growth is a consistently important part of the root cold response, even though its direction differs between the annual and the perennial species.

**Fig. 5.**
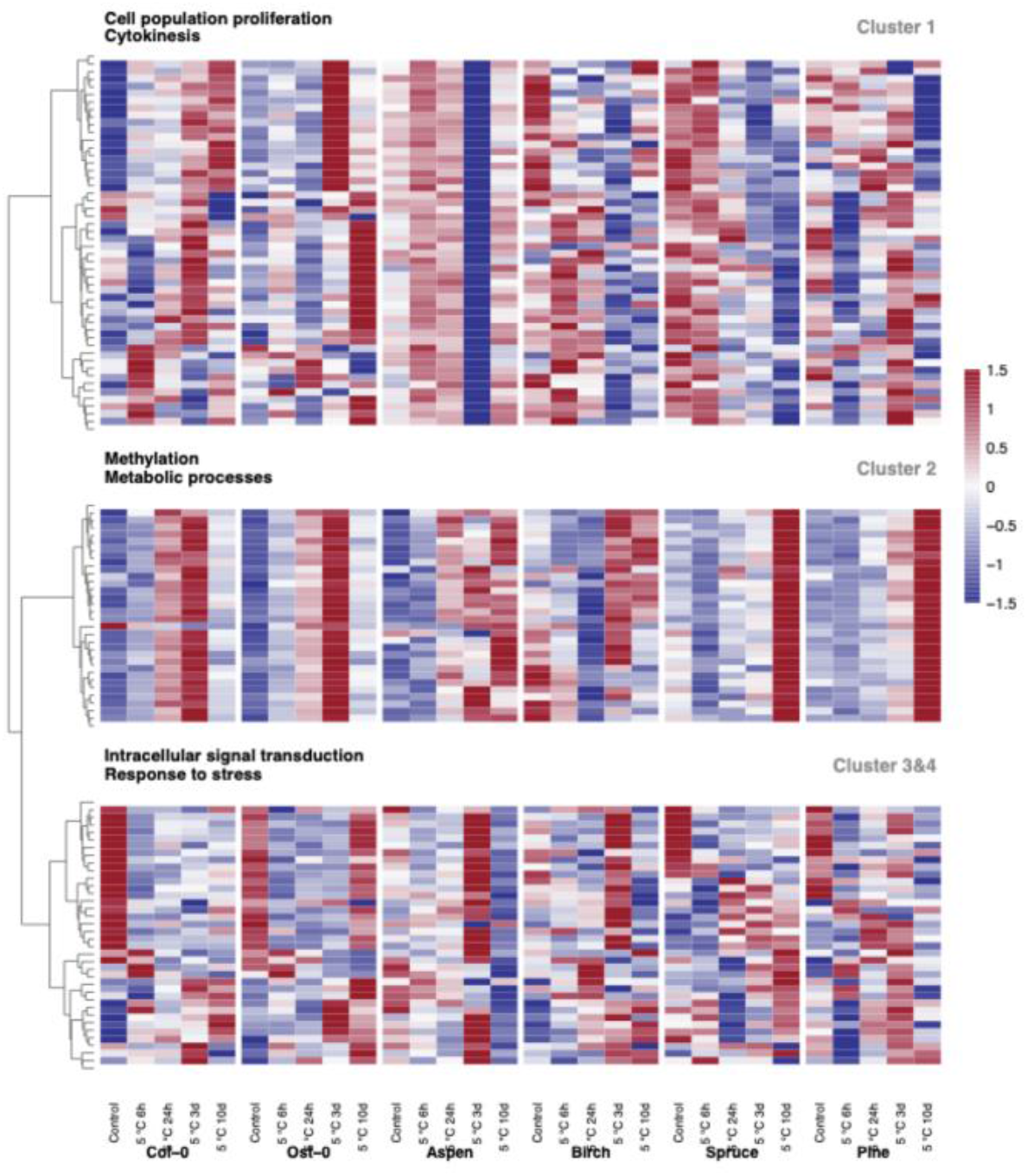
Heatmap of *Z*-score-scaled, VST-normalised median expression of genes within cliques. Each row is one orthogroup clique across all species; columns are ordered chronologically through the cold experiment.

We also checked the conserved core against a dataset independent of the expression data generated here, though itself derived in part from public stress-response experiments. If the clique clusters reflect real co-regulation, the Arabidopsis genes within a cluster should also be co-expressed in other studies. Using STRING, a public co-expression database assembled from data we did not use, we found that co-expression links fell within our clusters 3.4-fold more often than expected by chance (P < 0.001; Fig. S5b). This indicates the clusters capture genuine co-regulation rather than an artefact of our pipeline.

To identify candidate regulators of this conserved core, we examined eight cliques composed of transcription factors (Fig. 6; in one, the bZIP61 clique, the birch member is not annotated as a TF). These spanned seven families: LATERAL ORGAN BOUNDARIES DOMAIN (LBD), basic leucine zipper (bZIP), MYB, NAC, WRKY, heat-shock factor (HSF) and GARP (here the KANADI-family member KAN3). Several have established roles in cold-stress signalling (Zhou et al. 2011; Dong et al. 2021; Mei et al. 2023). For two of the eight cliques (NAC and WRKY), the member recovered in Ost-0 differed from that in Col-0, so in these cases the conserved relationship is carried by closely related paralogues within the orthogroup rather than by a single identical gene in every species. Notably, these TFs shared co-expression neighbourhoods across all species despite showing variable, and generally modest, changes in their own expression, identifying a conserved co-expression core whose species-specific fine-tuning may underlie the contrasting timing seen among the clique clusters.

**Fig. 6.**
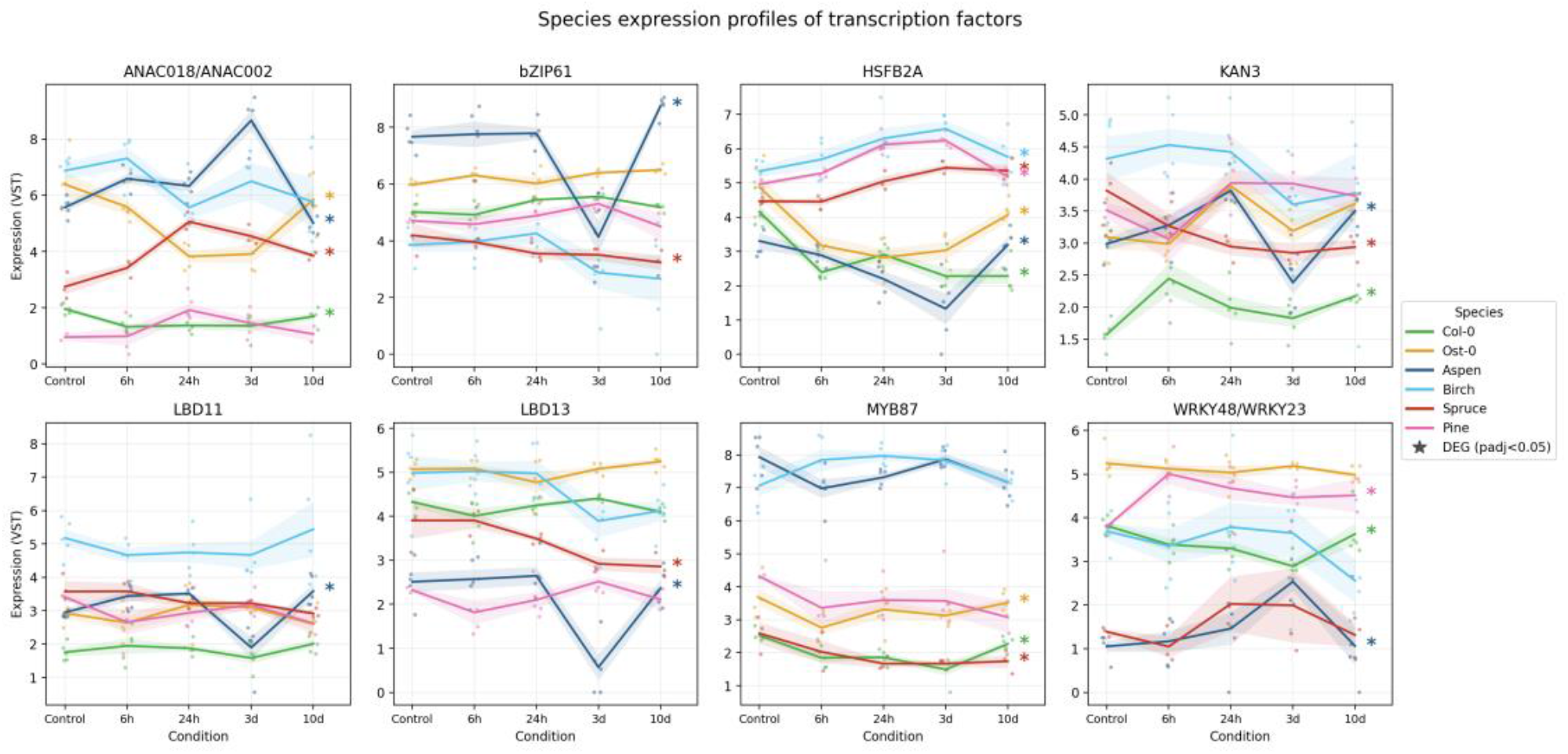
Species expression profiles of transcription factors within each all-TF clique. Each panel shows one clique of co-expressolog transcription factors, labelled by the Arabidopsis (Col-0) member; for the NAC and WRKY cliques the Ost-0 member differed from Col-0 (ANAC018/ANAC002 and WRKY48/WRKY23). Lines show mean VST-normalised expression for each species across the cold time course (control, 6 h, 24 h, 3 d, 10 d); individual replicates are shown as points and the shaded band denotes ± 1 standard error (n = 2–5 replicates per species and timepoint). Asterisks mark genes differentially expressed in that species during cold (DESeq2, adjusted P < 0.05).

## Discussion

Few comparative analyses of the root cold-stress transcriptome have spanned both angiosperm and gymnosperm trees alongside an herbaceous reference, and fewer still have used comparative co-expression to separate a conserved regulatory core from the largely divergent, species-specific differential-expression response of roots. Conserved cold-responsive transcription-factor orthogroups have recently been defined and, in some cases, functionally validated in angiosperms (Jia et al. 2026), against a backdrop of repeated, convergent evolution of cold adaptation across flowering plants (Wang et al. 2025b); our contribution is to ask whether such conservation extends across the angiosperm– gymnosperm divide and to recover it through co-expression rather than differential-expression overlap, in roots. Our central result is that these two views of the same data are not in conflict but are complementary: which genes are differentially expressed, and when, is strongly species specific, yet beneath this divergence lies a core of conserved co-expression relationships shared across more than 300 million years of evolution. This reconciles the reported poor transferability of cold responses with the conserved cold-acclimation programme: what is conserved over this depth of time is the coordination of expression, consistent with shared regulation, rather than the timing or magnitude of individual gene responses. Cold tolerance was, moreover, acquired largely independently in the two lineages: conifer freezing resistance is ancient and long-established (Sakai 1983), whereas many cold-adapted angiosperm lineages arose much more recently during Cenozoic cooling (Hagen et al. 2019), and their common ancestor is unlikely to have been under selection for cold. A cold-specific regulatory module would therefore not be expected to be shared across this divide, and the conserved core we recover is instead more general, comprising growth, metabolism and stress-signalling genes that each lineage appears to have recruited independently into its cold response.

### The differential-expression response of roots is largely lineage specific

The molecular cold response of roots has been studied far less than that of shoots, and existing work is dominated by herbaceous angiosperms (Moliterni et al. 2015; Zhao et al. 2021; Vergara et al. 2022). We found extensive divergence in the set of cold-responsive genes among species (Fig. 1), and because gene content was largely shared, this divergence reflects differential regulation of a common repertoire rather than differences in gene complement. Responses also differed markedly in timing, with Arabidopsis responding fastest and the trees more slowly (Fig. 2), and genes with shared temporal trajectories were generally not orthologous (Fig. 3). Divergence extended even to the two Arabidopsis ecotypes, consistent with the proposal that genetic variation in stress-responsive expression contributes to local climatic adaptation (Lasky et al. 2014); comparable among-population divergence in the autumn cold-acclimation transcriptome has been documented within conifers (Holliday et al. 2008).

These observations are congruent with a broad body of comparative work. The same pattern, conserved gene sequences but largely species-specific cold-responsive expression, has been reported for above-ground tissues of Norway spruce and Arabidopsis (Vergara et al. 2022), for drought responses of Norway spruce and Arabidopsis (Haas et al. 2021), for the temperate grass subfamily Pooideae (Schubert et al. 2019), for Arctic Brassicaceae (Birkeland et al. 2020; Birkeland et al. 2022).Our data extend this conclusion to roots and, importantly, across the gymnosperm–angiosperm divide.

A recurring interpretive difficulty is that differential expression under stress conflates regulated, acclimatory change with secondary or downstream change that need not be functional, and a substantial fraction of stress-induced transcriptional change may represent the latter. Comparative co-expression offers a principled way to distinguish the two, because the conservation of co-expression neighbourhoods depends on the persistence of functional regulatory relationships rather than on the response magnitude of any individual gene. Co-expression conservation has been shown to recover deeply conserved, functionally constrained gene programmes across kingdoms (Crow et al. 2022), and the architecture of regulatory networks itself shapes which genes vary and which are buffered (Chalancon et al. 2012). In *Populus*, genes occupying central, highly connected positions within the co-expression network experience stronger purifying selection on both coding and regulatory sequence than peripheral genes (Mähler et al. 2017), directly linking network architecture to selective constraint. Across more distantly related plants, the gene families most repeatedly involved in local adaptation to climate likewise occupy more central and pleiotropic positions in co-expression networks (Whiting et al. 2024), suggesting that network position shapes not only selective constraint but also which orthologues are repeatedly available for adaptation. This framework is consistent with the finding that genes with consistent stress-responsive expression are under stronger purifying selection than genes whose responses vary among genotypes, the latter being preferentially associated with local adaptation (Lasky et al. 2014). A comparable decoupling is apparent within conifers: in a comparison of lodgepole pine and interior spruce, genes diverging more rapidly in sequence also diverged more in expression, a relationship shaped primarily by negative rather than positive selection (Hodgins et al. 2016). Furthermore, a parallel study of Norway spruce and Scots pine roots and needles under cold and drought (reusing the conifer cold-root samples analysed here) found that genes with broader co-expression conservation are under stronger purifying selection on their coding sequences (a lower ratio of non-synonymous to synonymous substitutions, dN/dS) and, within Norway spruce populations, a lower ratio of non-synonymous to synonymous polymorphisms (pN/pS) (van Zalen et al. 2026), providing orthogonal, sequence-level evidence that conserved co-expression marks functional constraint. Viewed this way, the divergent differential-expression response we observe is expected, and the conserved co-expressologs represent the regulated core most likely to be functionally important.

### A conserved core centred on growth regulation

The strength of comparative co-expression is that conservation does not require shared expression dynamics, only shared co-expressed neighbourhoods (Netotea et al. 2014; Ovens et al. 2021; Crow et al. 2022), and this allowed us to recover a conserved core despite the contrasting timing of responses among species. Within this conserved set, 256 orthogroups had co-expressolog pairs present in every pairwise species comparison, of which 116 formed fully interconnected cliques (Fig. 5), the most stringent class of conserved relationship. The largest conserved cluster concerned the regulation of cell division and root growth, and showed a clear directional contrast: cold induced these genes in Arabidopsis but repressed them in the trees (Fig. 5, Cluster 1). One possible interpretation is that this reflects divergent life-history strategies — an annual maintaining root growth to exploit transient resources, versus perennials curtailing growth in preparation for winter (Singh et al. 2017); however, Arabidopsis differs from the trees not only in life history but also in growth form, in being grown from seed rather than as an established plant, and in husbandry (see Limitations), so life history cannot be isolated as the cause of this contrast here.

A gibberellin (GA)-related signature provides a concrete worked example of this conserved-but-redirected growth response, and gibberellin-related GO terms were enriched among the broadleaf–conifer (BC) co-expressologs in aspen and Norway spruce, and among the conserved core in Arabidopsis (Table S2). In Norway spruce, genes encoding gibberellin-biosynthetic enzymes that responded significantly to cold were predominantly down-regulated, consistent with a reduced pool of bioactive gibberellin that would stabilise the growth-repressing DELLA proteins and restrain growth; this trend was weaker and less consistent in the broadleaf trees, and we did not observe a coordinated up-regulation of the gibberellin-deactivating GA2-oxidases. Reduced GA signalling is an established route to growth restriction under cold and other abiotic stresses (Colebrook et al. 2014; Shu et al. 2018), and the transcriptional control of GA metabolism is a recognised point of regulation in this response; in Arabidopsi*s*, the membrane-bound NAC factor GIBBERELLIN SUPPRESSING FACTOR is processed and relocates to the nucleus under cold, where it activates *GA2ox* genes and suppresses GA biosynthesis (Chen et al. 2019). In woody species, inhibition of GA biosynthesis promotes growth cessation, bud set and cold acclimation (Mølmann et al. 2005; Welling & Palva 2006; Chang et al. 2021). Our data indicate that this GA-mediated growth restriction extends to the roots of boreal trees, contributing to the growth cessation that distinguishes the perennial trees from the herbaceous annual. As the GA–GID1–DELLA perception and growth-repression module was assembled in the ancestor of the vascular plants (Hernández-García et al. 2021) and is functional in conifers (Du et al. 2017), this shared growth-cessation signature is most parsimoniously interpreted as independent redeployment of a deeply conserved pathway rather than convergent evolution, with lineage-specific tuning of this ancestral module. A complementary comparison of Norway spruce and Scots pine spanning cold and drought in both needles and roots likewise places growth control within the conserved stress co-expression core (van Zalen et al. 2026), consistent with the enrichment of gibberellin-related functions among the conserved co-expressologs. Similarly, a comparable link between contrasting GA-mediated transcriptional responses and divergent cold acclimation has recently been reported between two diploid cotton species (Wang et al. 2025a), reinforcing GA as a recurring axis of cold-response divergence.

### Conserved transcription factors as candidate regulators

The TF cliques (Fig. 6) provide candidate regulators of the conserved core, several of which have known roles in cold or abiotic stress. The NAC factor ATAF1 (ANAC002), a drought- and abscisic-acid-induced regulator of stress-responsive genes (Lu et al. 2007) that is also induced by jasmonate and mediates responses to both abiotic and biotic stress (Wu et al. 2009), was up-regulated in aspen and Norway spruce but was down-regulated in both Arabidopsis ecotypes, and the enrichment of abscisic-acid terms in the tree co-expressologs (BC; Fig. 4) is consistent with a role for this factor in the root cold response. The R2R3-MYB factor MYB87 (Dubos et al. 2010) showed no clear cold induction in Arabidopsis but was constitutively high in aspen and birch. One of the conserved co-expressologs was a WRKY transcription factor. Although WRKY factors are typically associated with abscisic-acidmediated stress signalling (Zhang et al. 2016; Luo et al. 2017), the Arabidopsis member of this clique, *WRKY23*, instead functions in auxin-dependent root development through local control of flavonol biosynthesis (Grunewald et al. 2012), suggesting that this conserved module could link developmental and environmental signalling under cold. Notably, this clique was conserved across the angiosperm and conifer species, with the Scots pine orthologue among the cold-induced members. The heat-shock factor *HSFB2A*, a class B heat-shock factor characterised in Arabidopsis gametophyte development and heat-stress regulation (Wunderlich et al. 2014), was up-regulated in birch, Norway spruce and Scots pine at 3 days. By contrast, the less well-characterised LBD factors *LBD11* and *LBD13* (Cho et al. 2019; Dang et al. 2023), and *bZIP61* and *KAN3*, showed variable and generally modest responses without a consistent direction across species. That these regulators are recovered as co-expressologs despite small expression changes is itself informative: it indicates a conserved co-expression core whose outputs are tuned species-specifically, and it underscores that the bulk of each species’ large-scale transcriptional response lies outside this conserved core and is controlled by species-specific regulation.

### The canonical ICE1–CBF–COR pathway is not part of the conserved root core

Studies of above-ground cold acclimation in herbaceous angiosperms have placed the ICE1– CBF–COR module at the centre of the cold response (Stockinger et al. 1997; Liu et al. 1998; Chinnusamy et al. 2003; Qian et al. 2024), yet neither *ICE1* nor the *CBFs* were themselves members of a conserved co-expressolog clique. This is consistent with the evolutionary history of the pathway, but also with the limited power to detect small, lineage-specific gene families and with uncertainty in *CBF* orthology assignment across gymnosperms (Fig. S6).

The CBF/DREB1 factors arose by duplication in ancient angiosperms and expanded independently in eudicots and monocots (Nie et al. 2022), consistent with the widespread retention of duplicated regulatory genes in angiosperms following polyploidy (Wu et al. 2020). Even among the trees studied here, *CBF* regulation is elaborate: birch carries multiple cold-responsive *CBF* paralogues that are differentially regulated between growing and dormant tissues, with their induction delayed in the dormant state (Welling & Palva 2008). Consistent with this, the conifers had far fewer genes co-expressed with the *CBFs* than the angiosperms (Fig. S6), and *CBF* expression was highly variable among species. Reflecting this, Vergara et al. (2022) found only a single, weakly cold-induced *CBF1*/*CBF3* orthologue, whose expression fell below the threshold for DEG/COR classification and was detected in roots but not needles, indicating limited conservation of the *CBF* node specifically. *ICE1*, by contrast, emerged in that same study as the most highly connected hub of the Norway spruce cold-regulatory network, underscoring that its absence from our conserved co-expressologs here does not reflect a lack of regulatory importance in individual species, but rather that its co-expression neighbourhood is not shared across the wider set of species compared in this study. Thus, the lineage-restricted evolution of ICE1–CBF–COR offers a plausible explanation for why a pathway so central to angiosperm cold acclimation does not form part of the response conserved across all species in our study, and reinforces the value of co-expression-based approaches that do not presuppose a particular regulatory module.

### Limitations and future directions

Our analyses are based on expression and orthology and do not incorporate DNA-sequence-level information; with the phylogenetic sampling used here we therefore cannot formally exclude convergent evolution as an alternative to common descent for some conserved responses, and we refer to conserved responses in this sense. Husbandry also necessarily differed between the herbaceous reference and the trees (including plant age, pot size and photoperiod), so absolute differences in the timing of the response between Arabidopsis and the trees should be interpreted with corresponding caution; this does not, however, affect the within-species co-expression relationships on which the conserved-core analysis depends. Fine roots (under 2 mm) were used as the comparable unit across species, although root-system architecture and developmental stage differed between the herbaceous Arabidopsis and the woody seedlings, and exact tissue equivalence cannot be guaranteed. Our comparisons are also framed at the level of orthogroups and do not capture intraspecific presence–absence or copy-number variation in gene content, which can itself shape stress-response repertoires; resolving its contribution will require pangenome-level sampling within each species. The conifer-specific co-expressologs returned no functional enrichment against the Norway spruce annotation used here; tested instead against the Scots pine annotation, this set is enriched for ubiquitin–proteasome terms (led by SCF-dependent proteasomal ubiquitin-dependent catabolism), so the apparent absence is annotation-limited, and improved, harmonised conifer annotation would allow this lineage-specific component to be interpreted more fully. Functional (GO) enrichment for cross-species sets relied on the best-annotated genome in each comparison (Arabidopsis, or aspen or Norway spruce where Arabidopsis was absent), which may bias the recovered terms towards functions already characterised in Arabidopsis. The dataset also contains extensive species- and lineage-specific responses that we have not explored here, and these constitute a resource for future work. Control samples were harvested at a single reference timepoint rather than being time-matched to each cold timepoint; over the 10-day treatment, ontogenetic drift is expected to be negligible in the perennial tree roots but may contribute to the 10-day Arabidopsis contrast, and the later Arabidopsis timepoints should be interpreted accordingly. Replication was also unequal across species (15–25 samples per co-expression network) due to stochastic loss of some replicates at various stages of the study; because a |log2FC| ≥ 1 significance threshold is crossed more readily in species with more replicates and lower dispersion, some of the apparent between-species differences in DEG number may reflect this imbalance rather than biology. Annotation completeness also differs among the genomes, and this asymmetry may inflate the Arabidopsis-containing pairwise co-expressolog sets relative to the conifers; the conserved core, however, requires a co-expressolog in all fifteen pairwise comparisons and is therefore bounded by the least-completely-annotated (conifer) genomes, so it is relatively insulated from this effect. Natural extensions include broadening the phylogenetic sampling, integrating population-genetic signatures of selection to link regulatory divergence to local adaptation, and functionally validating candidate regulators such as ATAF1 and the conserved MYB and WRKY factors in tree systems.

## Conclusion

We present evidence for a conserved set of genes regulated in response to cold in the roots of boreal trees and a herbaceous reference, encompassing growth regulation, metabolism and stress signalling, in which a GA-associated growth-cessation signature is prominent in the trees. The timing and, for growth-related genes, the direction of this conserved response vary among species, and the bulk of each species’ transcriptional response lies outside the conserved core and is species specific. Two conclusions follow. First, comparative co-expression recovers conserved regulation that is invisible to differential-expression analysis alone, and is therefore a powerful approach for cross-species comparison when response dynamics differ. Second, and of practical importance, the set of genes differentially expressed under cold in one species is a poor direct guide to that in another, including between ecotypes of a single species, underscoring the need for species-level study when translating cold-response biology among boreal trees.

## Author contributions

N.R.S. and V.H. conceived and designed the study. T.A., E.v.Z., C.C. and V.K. performed the experiments and generated the RNA-sequencing data. T.A., E.v.Z., A.V., E.D.C. and T.R.H. carried out the bioinformatic and statistical analyses. T.A. and E.v.Z. wrote the first draft of the manuscript. N.R.S., V.H. and T.R.H. supervised the work and acquired funding. All authors contributed to the interpretation of the results, revised the manuscript critically and approved the final version.

## Data availability

The raw RNA-sequencing reads are available in the European Nucleotide Archive (ENA) under umbrella BioProject PRJEB104158, which groups the component projects PRJEB104117 (Arabidopsis), PRJEB104118 (birch), PRJEB104119 (aspen), PRJEB104120 (Scots pine) and PRJEB26918 (Norway spruce, fine roots). The root cold-stress samples of the two conifers are shared with the companion study of van Zalen et al. (2026), in which they are also analysed: the Scots pine samples are first reported here, and the Norway spruce samples were originally generated by Vergara et al. (2022) and are reanalysed in both studies. Processed gene-level expression matrices, co-expression networks and the analysis scripts supporting the findings are available from FigShare (DOI:10.17044/scilifelab.32747586) and GitHub (https://github.com/natstreet/RootColdProject; DOI:10.5281/zenodo.21628010). The genome and transcriptome assemblies used for read quantification are publicly available as described in the Materials and methods.

## Acknowledgements

We thank colleagues at the Umeå Plant Science Centre (UPSC) for discussion. We specifically thank the UPSC bioinformatics platform and the greenhouse support team.

## Supporting Information

The following Supporting Information is available for this article:

**Fig. S1** Correlation-based identification of conserved cold-responsive transcriptional profiles across species. Pairwise Pearson correlation matrix of mean expression profiles from TF-DEG clusters across six species (36 total cluster profiles: 6 species × 6 clusters per species). Colour intensity represents correlation strength and direction (blue: positive; red: negative; white: no correlation). Square size indicates absolute correlation magnitude, with larger squares denoting stronger correlations. Clusters showing high positive correlations (large blue squares, r > 0.7) were manually grouped into 12 superclusters (SC1–SC12), representing conserved cold-responsive expression patterns. superclusters vary in phylogenetic breadth, from broadly conserved patterns present across all species to lineage-specific responses.

**Fig. S2** Transcription factor (TF) family composition across plant species in superclusters. Each bar represents the relative contribution of five representative TF families (ERF, MYB, NAC, bHLH and WRKY), with all other families grouped as “Others”. Panels correspond to superclusters SC_I–SC_IV, and the number of transcription factors per species is shown above each bar.

**Fig. S3** Bar chart of the number of genes per species classed as co-expressologs, and as differentially expressed (DE) co-expressologs.

**Fig. S4** UpSet plot of orthogroup overlaps between co-expressologs. Horizontal bars show the number of overlapping orthogroups within the co-expressologs of the pairwise comparisons. Vertical bars show the overlap of orthogroups within the co-expressologs across all comparisons. All co-expressologs within this plot are p < 0.1.

**Fig. S5** Robustness of the conserved co-expression core. (a) Permutation test: across 5,000 permutations that randomised co-expressolog assignments within each pairwise comparison, no orthogroup was recovered as a co-expressolog across all 15 comparisons (null distribution), versus the observed conserved core (red line; P < 0.0002). (b) Cluster coherence in independent data: among the conserved-core Arabidopsis genes, co-expression edges in STRING fell within clique clusters (red line, observed = 692) 3.4-fold more often than when the same genes were randomly assigned to clusters (null mean approximately 203; P < 0.001).

**Fig. S6** Expression profile of co-expressologs (P-value < 0.1) directly linked to Arabidopsis CBFs (CBF1: AT4G25490; CBF2: AT4G25470; CBF3: AT4G25480). Expression is plotted as median VST values.

**Table S1** GO enrichment of the 270 commonly differentially expressed orthogroups (topGO classic Fisher, biological process).

**Table S2** GO enrichment of conserved co-expressolog sets (ABC core; AB and BC sets), per species.

**Table S3** GO enrichment of the conserved-core clique clusters, per species (manuscript cluster numbering).

**Table S4** Gene membership of superclusters SC_I–SC_IV, per species/ecotype. One worksheet per supercluster; each column lists the differentially expressed genes assigned to that supercluster in the given species, with Col-0 and Ost-0 shown separately. A final worksheet (TF_families) lists the transcription factors among the members and the family to which each belongs (680 unique loci; because Col-0 and Ost-0 share Arabidopsis gene identifiers, ecotype-specific transcription-factor membership is resolved in the per-super-cluster worksheets).

