## Supplementary figures and images for "Comparative co-expression reveals a regulatory core shared by angiosperm and conifer roots under cold"

### Fig. S1

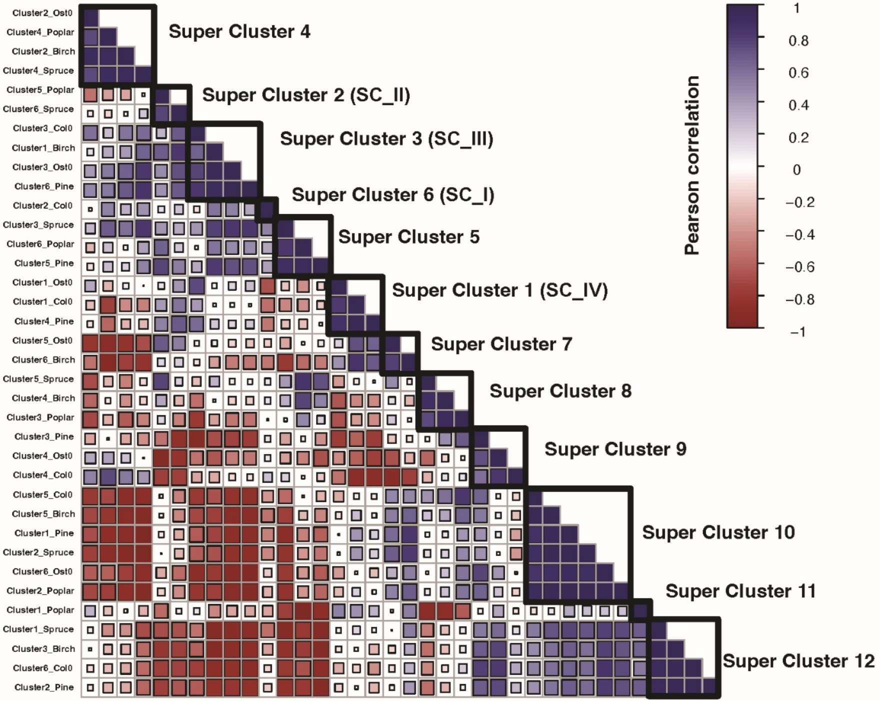

### Fig. S2

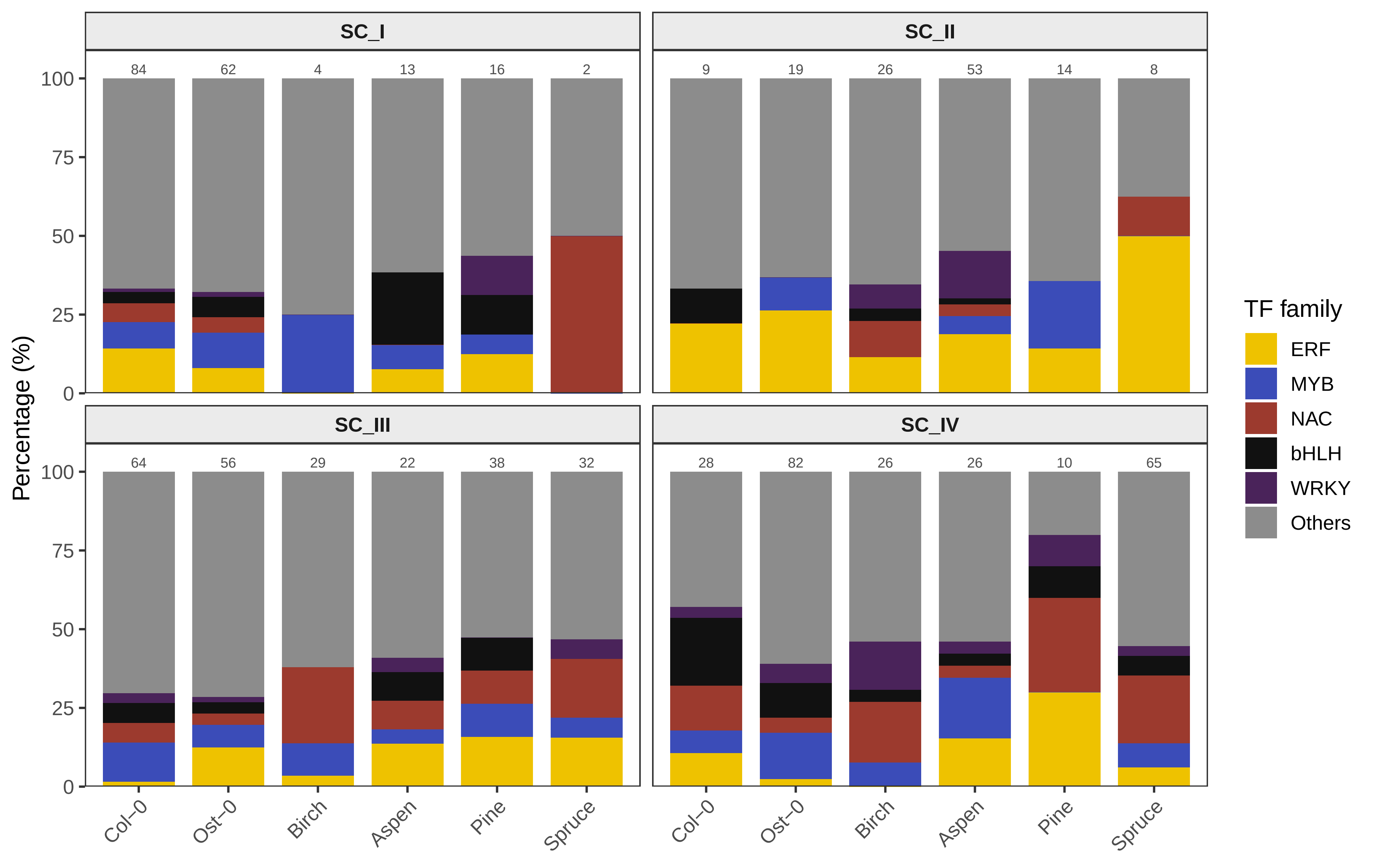

### Fig. S3

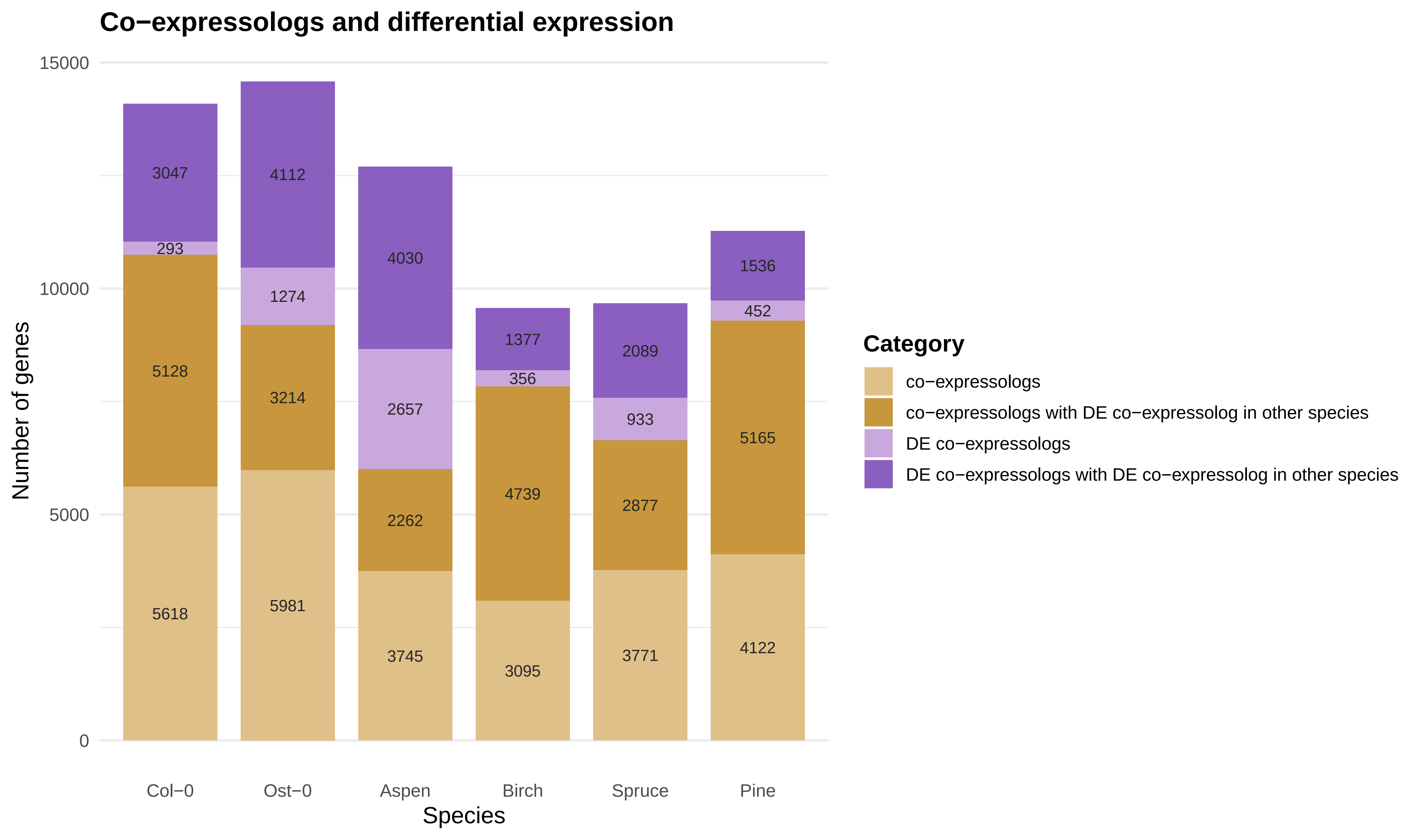

### Fig. S4

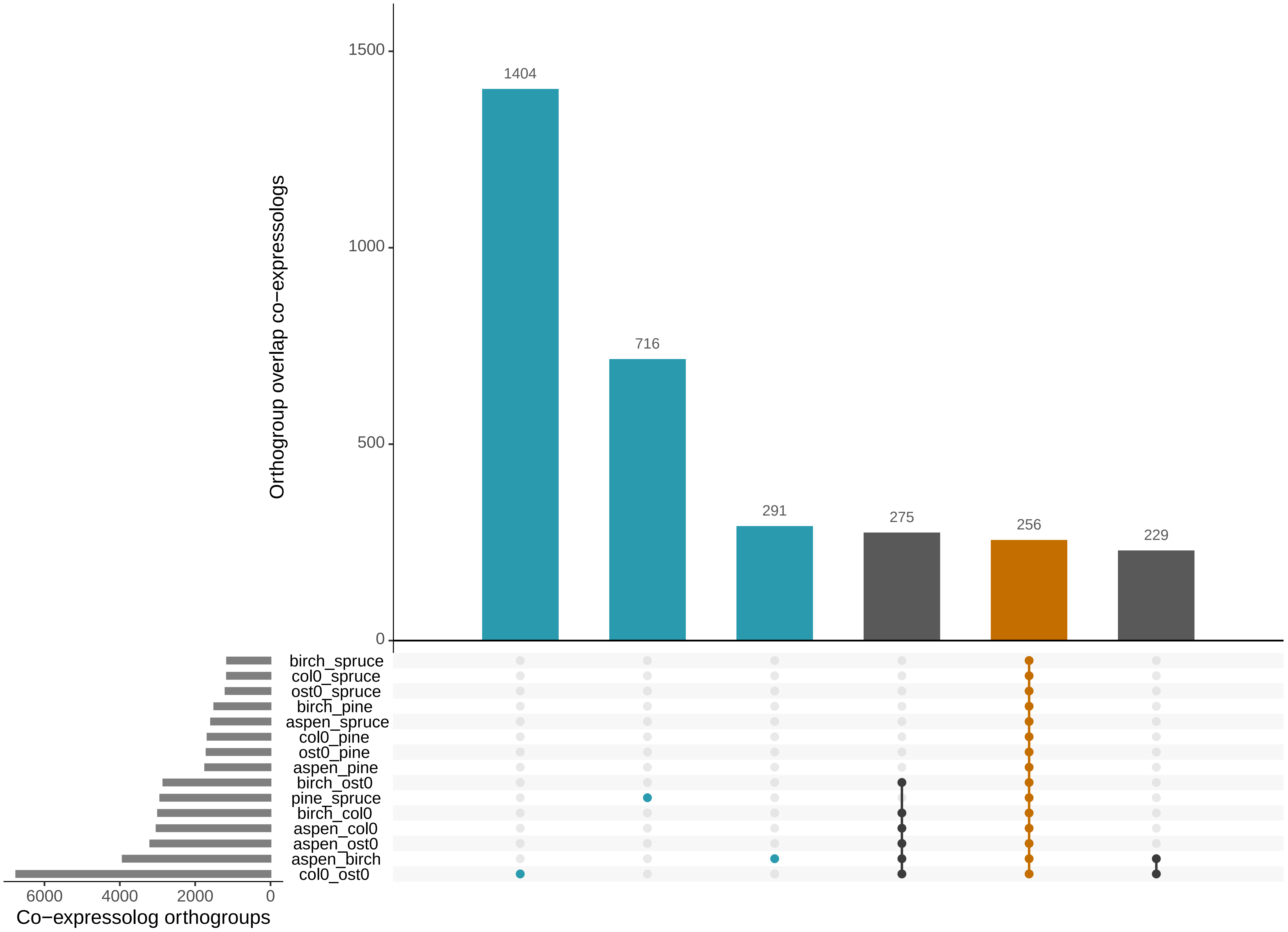

### Fig. S5

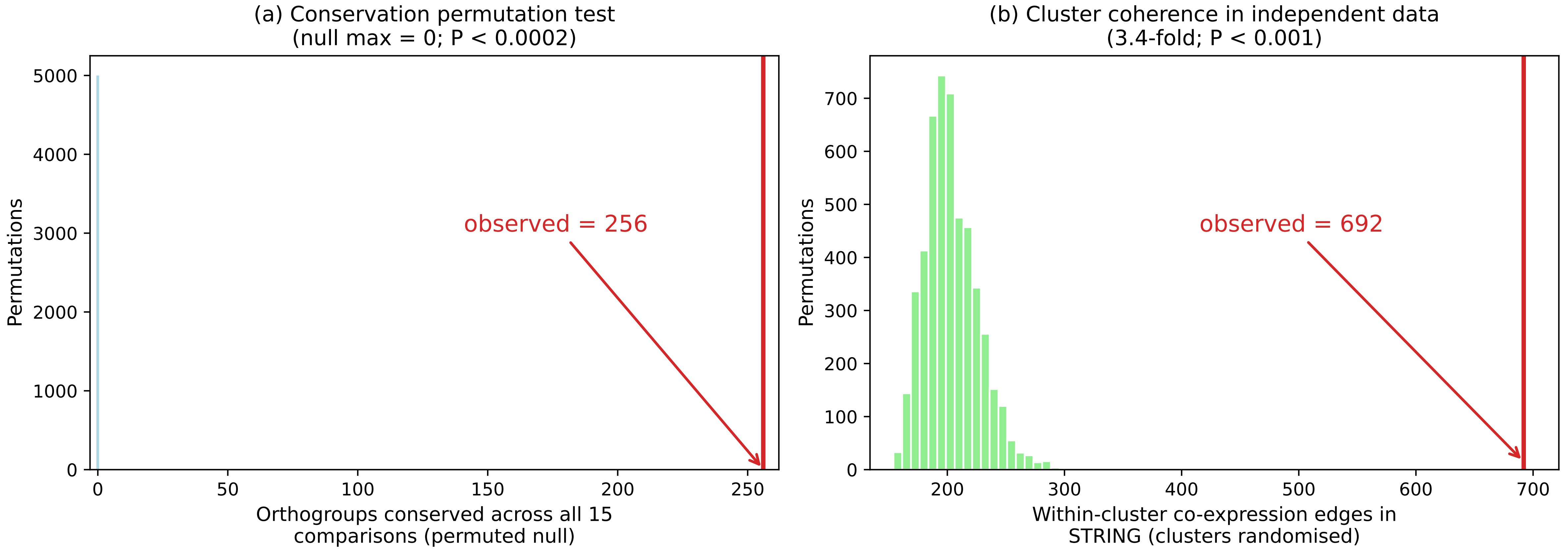

### Fig. S6

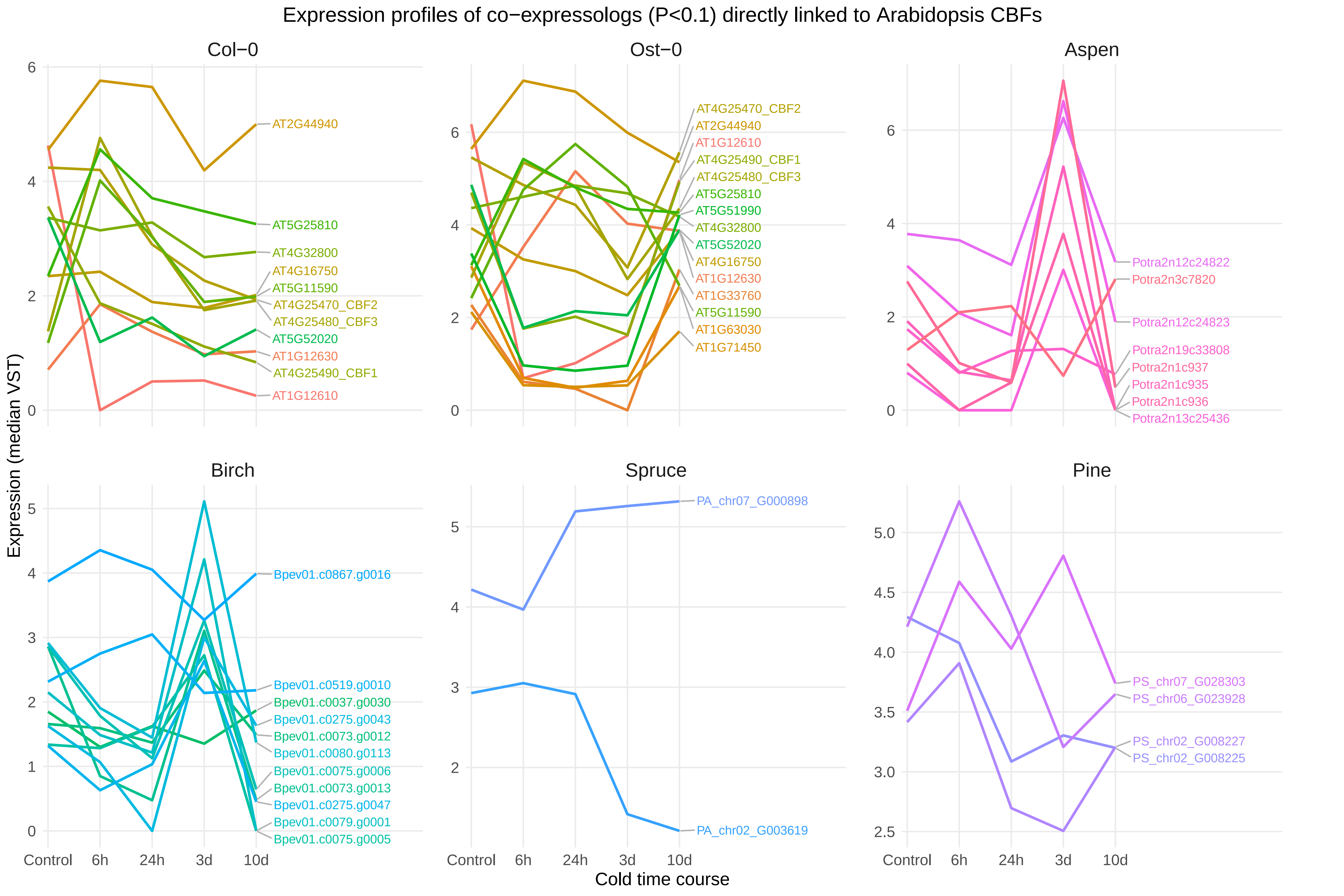
